# Biomechanical stress, inflammatory change and atheromatous plaque detachment – the role of pulse pressure, arterial wall stiffness and vessel bending : an in-vitro experimental study with some clinical implications

**DOI:** 10.1101/2025.07.03.662946

**Authors:** Thomas K Day, Ellie Newman

## Abstract

The dynamic forces contributing to arteriosclerotic plaque separation were studied *in-vitro* using fresh porcine arteries and artificial plaques of different thicknesses mounted on thin ferro-magnetic strips. The arteries were subjected to pulsatile perfusion using an apparatus in which dynamic and rheological variables could be varied and controlled. The “plaques” were held on the inner wall of the artery by the force exerted by an externally-mounted electromagnet. The pressure retaining the artificial plaque to the wall could be regulated by varying the power supplied to the magnet. The separation of the plaque from the vessel wall, characterised by the appearance of a cleft beneath it with each pulsation, was observed with M and B mode ultrasound. In this way the critical detachment force (CDF), acting on the plaque at the moment of first separation of the plaque, could be calculated. 10 arteries were used during the study, using multiple observations in single arteries to reduce inter-artery variation. The CDF was found to correlate closely with pulse pressure (r= 0.9, p<0.001). Wall stiffness was found to determine pulse pressure when other parameters were kept constant, irrespective of whether the stiffness was induced by raising mean arterial pressure or by stiffening the artery artificially with formalin. In comparing plaques producing approximately 20% and 40% occlusion there was roughly a doubling of CDF, but no further increase between 40% and 60% (t = 4.0, p<0.001). Longitudinal tortuous movement of the artery (arc bending) had a profound effect on CDF, increasing progressively with the angle of bend (Rs 1.0, p <0.001). In this model the main dynamic contributions to plaque stress are: (i) moderate stenosis in the presence of sustained flow; (ii) high pulse pressure, to which arterial wall stiffness in distributive arteries contributes (iii) arc bending. The clinical implications of the findings of this study are discussed, and it is proposed that recurrent dynamic wall stress, in addition to its direct effects on the plaque, may also play a part in the inflammatory determinants of plaque vulnerability.

## Introduction

Previous experiments studying the dynamic causes of transmural stress in healthy arteries subject to pulsatile flow in-vitro showed that pulse pressure and pressure waveform were the main determinants of the stress and that arterial wall stiffness played an important secondary role in affecting both pulse pressure and pulse pressure waveform. ^1^ It was postulated that the patchy distribution of atherosclerosis throughout the vascular tree and the role of known predisposing cardiovascular risk factors might be explained on the basis of localised foci of mechanical strain resulting from this stress. However these experiments were restricted to studying the stress arising across the smooth wall of a healthy vessel. The presence of atherosclerotic plaque on the inner lining of the vessel wall might be expected to affect the stress across the vessel wall and in particular the stress on the plaque. This is important because detachment of the plaque from the vessel wall underlies most cases of acute coronary syndrome and sudden cardiac death, and is implicated in some strokes, transient ischaemic attacks and distal embolic phenomena. ^2–5^ Several mechanisms are theoretically implicated: the Bernouilli effect, the torsional component of wall shear stress, and elastic compliance mismatch between plaque and vessel wall. The present study was designed to investigate the interplay of these elements using the previously-described in-vitro model, modified by the addition of an artificial “plaque” and employing a force-balancing technique to estimate the force acting on the plaque at the moment of its separation/rupture.

## Materials and methods

### Arteries and perfusion

Abdominal aorta specimens taken from freshly slaughtered adult Large White pigs were obtained from an abatoir conforming to European animal welfare standards (EC regulation 1099/2009) and were preserved in phosphate buffered saline at 4 deg C. The arteries were dissected from surrounding connective tissue, whilst being kept moist and the lumbar branch arteries were tied off with 2/0 silk at their origins. The segment used extended from the renal vessels to the bifurcation. The average length was 14.7cm and the mid-vessel diameter 18.3 (+/- 7)mm. The artery was mounted on the perfusion device previously described^1^ for delivering pulsatile flow under controlled pulse rate, waveform and pressure configurations. Figure 1 shows the arrangement of the artery in the test bath. Each artery was mounted on tubes at either end using a double turn of 2mm latex tube to hold it in place (Fig. 1) and submerged in the test bath up to the level of the upper surface in cool buffered saline. The artery was orientated so the lumbar branches were posterior, the flow going from proximal to distal. The magnet and the ultrasound probe impinged gently on the superior surface when the artery was filled. Once the artery was mounted, the tubes at either end were then moved apart to produce the required degree of longitudinal tension. The tubes could subsequently be approximated to study the effect of different degrees of arc bending.

**Figure 1.**
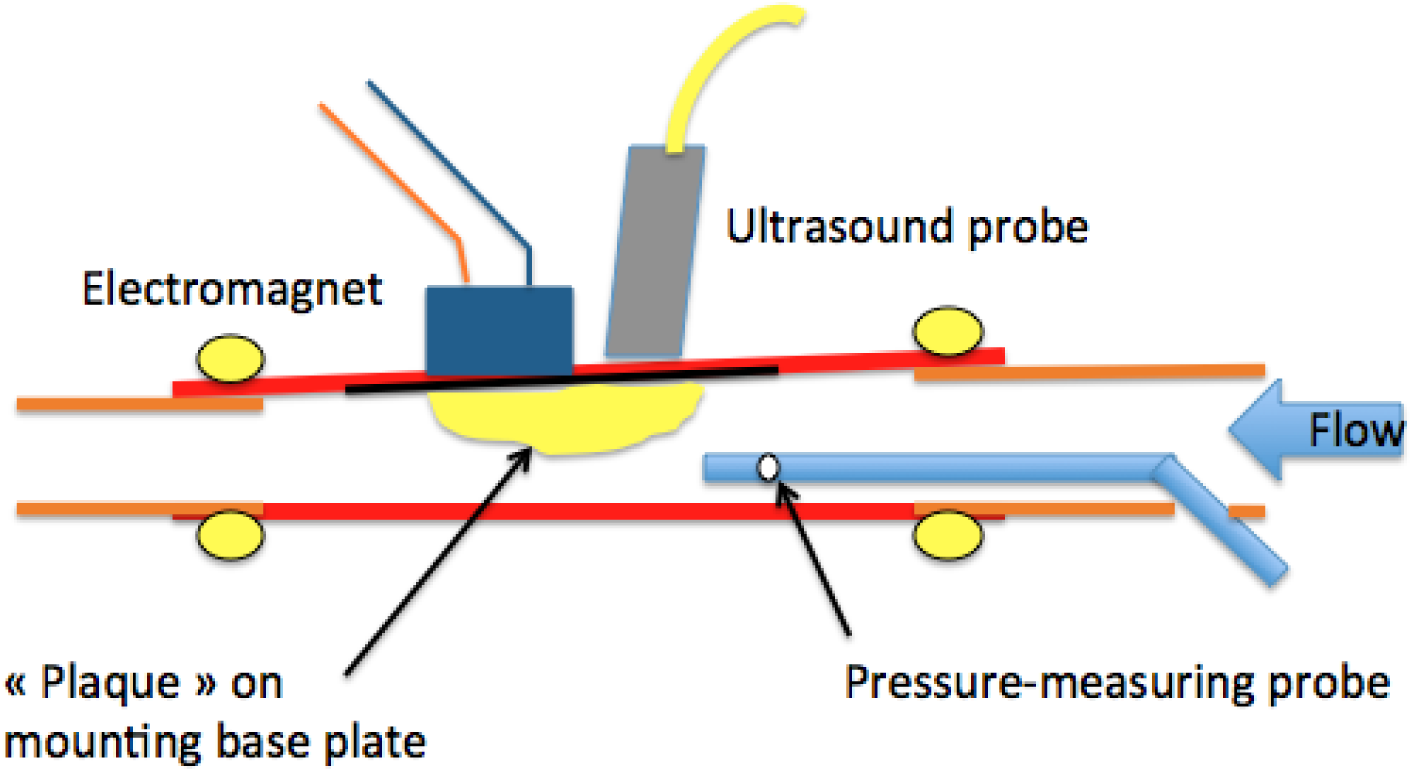
Experimental arrangement for measuring the force displacing the artificial plaque from the wall of the artery whilst observing the corresponding arterial pressure, pulse waveform, plaque separation and arterial wall movement using B/M mode ultrasound.

Pulsatile flow was induced by means of a cam-operated 60ml pump operated by one of four interchangeable cams of different profiles, each simulating a specific abdominal aortic pressure waveform. ^1^ The pulse waveform and volume load were determined by the profile of the cam. The cams were driven by a variable speed electric motor. The standard perfusate used was isotonic buffered saline, viscosity 1.0 M P, SG 1010. The effect of changing the viscosity and density of the perfusate was undertaken by comparing this solution with 40% sucrose with a viscosity of 7.4MPa.s, at 15 deg C, SG 1020 to simulate a hyper-viscosity state.

During test runs to investigate the effect of changing viscosity and density these solutions were alternated with buffered saline to provide multiple reading of CDF using a single artery as its own control. Arterial wall stiffness was adjusted by exposing the artery to 40% formaldehyde vapour for periods of up to 24 hours; hoops cut from the proximal end were used to assess the stiffening effect at 6-, 9- and 24-hours exposure, alternate hoops kept in cool buffered saline were employed to act as controls.

### Artificial Plaques

Simulated "plaques" were formed using pieces of moulded epoxy-resin mounted on thin flanges of mild steel 80x10x2mm, the weight of the resin being adjusted to give a total weight to the entire assembly of 16.0g. Three plaques were constructed with thicknesses of 3, 7 and 11mm. The force per unit area retaining the plaque to the arterial wall was estimated using a force balance technique, employing an electromagnet to retain the plaque on the inner arterial wall with a negative pressure that could be varied according to the power supplied to the electromagnet. The selected plaque was inserted into the lumen of the artery with its baseplate resting on the superior wall and was temporarily held in the chosen position using a small permanent magnet whilst the artery was mounted on the perfusion apparatus. Once mounted with the plaque in the chosen position, the plaque was held in place by the electromagnet and the permanent magnet removed. The effect of varying the thickness of the plaque on the CDF was estimated by replacing the plaques sequentially in the same artery in the same position so as to reduce the inter-artery variation due to dimensional changes and differences in elastic properties. A minimum of 5 runs was performed on each plaque/pulsatile pressure combination to determine the mean CDF.

### Electromagnet

A Heschen HS-P65 x 30 (Heschen Electric Co. Foshan, China) was employed. The vertical position of the magnet in relation to the artery was adjusted so that it contacted the superior wall of the artery without depressing it when the artery was filled but not yet subject to pulsatile flow. Current was applied using a regulated stabilised power source (UOHHBOE, Kefeixing Technology, Shenzen) and adjusted until the plaque was held firmly on the wall, its position being checked on the ultrasound. During each test run, measurement of the CDF was recorded at the moment when a cleavage plane beneath the plaque just started to appear during the diastolic phase of each pulsation. Each measurement of CDF was undertaken by slowly reducing the power supplied to the magnet until the cleft appeared. The power of the magnet was then increased a fraction until the plaque bedded down once more, and the test repeated, the mean value being recorded after 5 measurements.

After completion of each series of test runs, the permanent magnet was replaced in position to hold the plaque, the inferior surface of the artery was then split longitudinally, and the relationship between the power supplied to the magnet and the retention force exerted on the plaque was calibrated using a tensiometer connected to the plaque baseplate. The pull on the plaque was progressively increased until the tension peaked and then started to diminish. The peak tension was recorded as the critical force corresponding to each specific power setting, and the resulting graph used to estimate the CDFs these being recorded as force over unit area of the baseplate (Kpa).

### Ultrasound

Simultaneous B and M mode ultrasound was used to observe plaque separation and arterial wall dynamics during pulsatile perfusion. A 7.5 MHz probe was mounted just upstream of the magnet resting lightly on a small amount of contact jelly placed between the probe and the superior external arterial wall. The position of the probe was adjusted so as to observe the pulsatile movement of the posterior arterial wall and the position and degree of occlusion produced by the plaque.

Loss of image was found to be usually due to bubble formation in the perfusate, commonly caused by introduction of air if the level of fluid in the perfusate reservoir was allowed to fall too low, resolved by refilling the reservoir, running the circulating pump and manually squeezing the artery until the bubbles were expelled and a clear ultrasound image obtained.

### Runs and statistical methods

Observations in test arteries were undertaken in "runs" a run consisting of a series of tests each following a standard protocol in relation to changes in pulse rate and pulse pressure, fluid viscosity, percentage plaque stenosis and pulse waveform. The CDFs recorded during these runs was assumed to follow normal distribution. When assessing the effect of a single variable, such as pulse rate, multiple runs of data were obtained and harvested by groups to obtain observations on single-variable effects. When observations were made in respect to changes in plaque thickness, pulse waveform or perfusate viscosity, repeated observations in individual arteries were used to eliminate the noise produced by amalgamating the results from multiple arteries caused by individual differences in geometry and wall properties.

Comparison between groups was undertaken using the Kruskall-Wallis test for non-parametric variables, and correlation between non-parametric variables with Spearman’s rank correlation coefficient.

Arithmetical means with standard distribution was used in describing parametric variables.

## Results

### Pulse pressure, arterial wall stiffness and transmural stress

Figure 2 shows the relation between pulse pressure and the transmural stress acting on the plaque (r = 0.9, n=13, p<0.001) in a single fresh artery with a 7mm occluding plaque producing an approximately 17% occlusion at a diastolic pressure of 80+/- 10mmHg and a pulse rate between 55 and 65 bpm using a perfusate viscosity of 1.2cP

**Figure 2.**
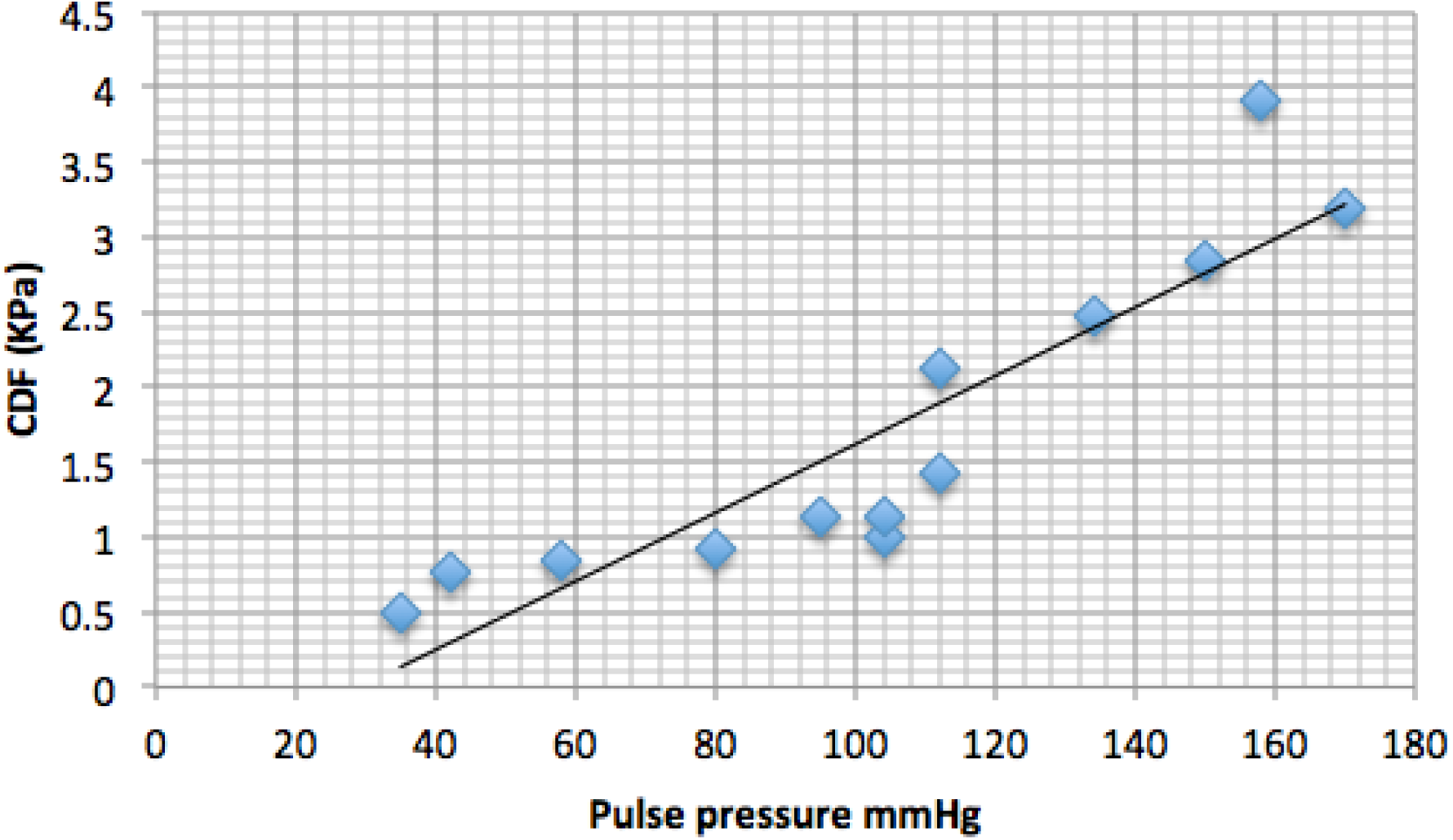
Close relation between the pressure acting on the plaque (critical detachment force, CDF) and the pulse pressure at the initiation of the sub- plaque cleavage plane.

This relation between pulse pressure and CDF was found to apply for plaques producing approximately 17%, and 40% occlusion irrespective of other variables.

Pulse pressure in this apparatus was associated with calculated peak systolic flow velocity: a pulse pressure of 50mmHg produced a calculated mean peak systolic flow velocity of 5.8cms (+/-1.2)cm/sec and that of 150mmHg a mean flow velocity of 8.9cms (+/-2.7)cm/sec.

Fig. 3 shows the relationship between mean arterial wall stiffness, pulse pressure and pulse waveform in a single typical vessel where the wall has been progressively stiffened by exposure to 40% formaldehyde over 24hrs. The stroke volume, cam profile and outflow resistance have been kept constant. As the vessel wall stiffens (hoop stretch Young’s modulus in KPa, is shown in the top line) the pulse pressure and consequently the transmural stress acting on the plaque rises, the upsweep gradient (dP/dT) increases and the systolic/diastolic transition becomes sharper. It is over this transition period that maximum stress is seen to occur across the arterial wall (fig, 4)).

**Figure 3.**
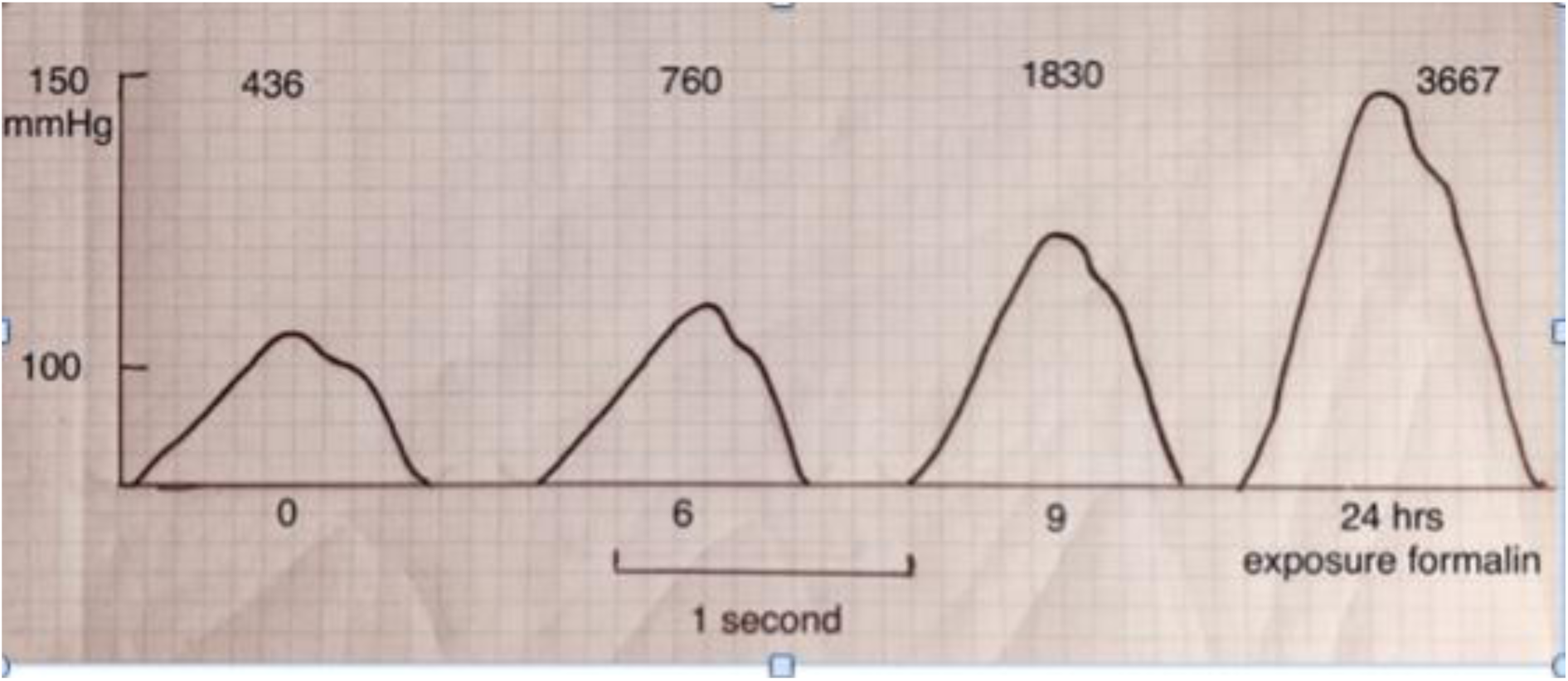
Under conditions of constant stroke volume and outflow resistance, reduction of elastic compliance in great vessels results in increase in pulse pressure, sharpened peak pressure waveform and consequently increased stress on the plaque. Here a pig’s aorta has been progressively stiffened by exposure to formalin vapour, simulating the effects of age, hypertension, disease, smoking and diabetes on elastic compliance. ^6,7^ The figures above the curves give the corresponding circumferential stretch elastic modulus corresponding to the degree of stiffness evoked.

### Interaction between arterial pressure and wall stiffness

Throughout the pressure cycle passive arterial wall stiffness responds in a non-linear fashion to instantaneous arterial pressure. Mean arterial pressure calculated as the geometric mean reflects the overall level of passive wall stiffness. Figure 4 illustrates the step-wise increase in wall resistance to stretch that occurs as the upper limit (for the pig ^8^) of physiological mean arterial pressure is reached.

**Figure 4.**
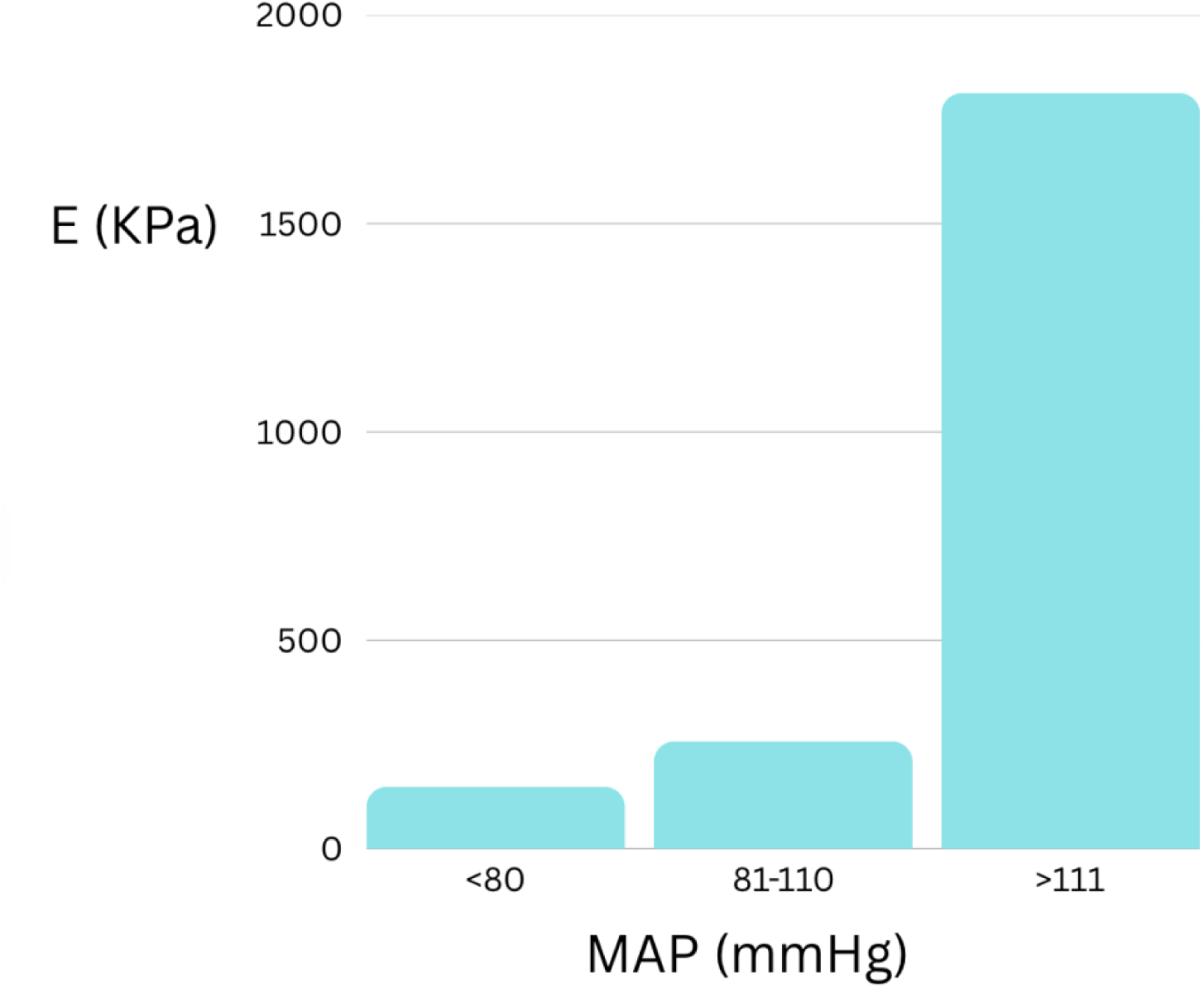
The effect of raising mean arterial pressure in healthy untreated vessels. When MAP increases above the physiological norm^8^ 81-110mm Hg, circumferential stretch modulus of elasticity increases about 7-fold reaching statistical significance (H= 24.5, p = <0.001). The dilatational response to increased pressure beyond this point becomes flat as the compliance limit of the artery is reached and the stress/strain ration decreases. This is accompanied by a rise in the pulse pressure and the distal transmission of sharp peaked high pressure pulse waves (fig.3). (n = 48 runs, 3 fresh arteries, pulse rate 60)

### Perfusate viscosity

Maintaining the machine settings for pulse waveform, diastolic pressure, volume load and pulse rate, use of a much denser and more viscous sucrose solution mimicking a hyper-viscosity state resulted in a modest rise in systolic and pulse pressure, but is also independently associated with a significant rise in CDF when controlled for pulse pressure close to or within the physiological range (P<0.001, t = 4.25, n= 13).

At pulse pressures greater than 111mmHg a trend was observed that fell short of statistical significance.

### Dynamic strain reactions on the vessel wall

Plaque detachment is normally seen to occur at systolic/diastolic transition and at both systolic and diastolic transitions when the mean arterial pressure is raised above the physiological range and the artery stiffens. The timing of strain separation of the intima and the two elastic lamina of the aorta is visualised in high-definition ultrasound (Plate 1).

It might be expected that steep pressure gradients would be associated with increased stress on the plaque in view of the known viscoelastic qualities of the arterial wall. However, this was not demonstrated in the current model. The combination of high pulse pressure and rapid pulse rate did lead to an augmentation of upsweep pressure gradient (dP/dT) irrespective of MAP (Spearman’s R = 0.72, p = 0.002, n= 15), but dP/dT was not independently correlated with CDF when controlled for pulse rate and arterial pressure.

**Plate 1.**
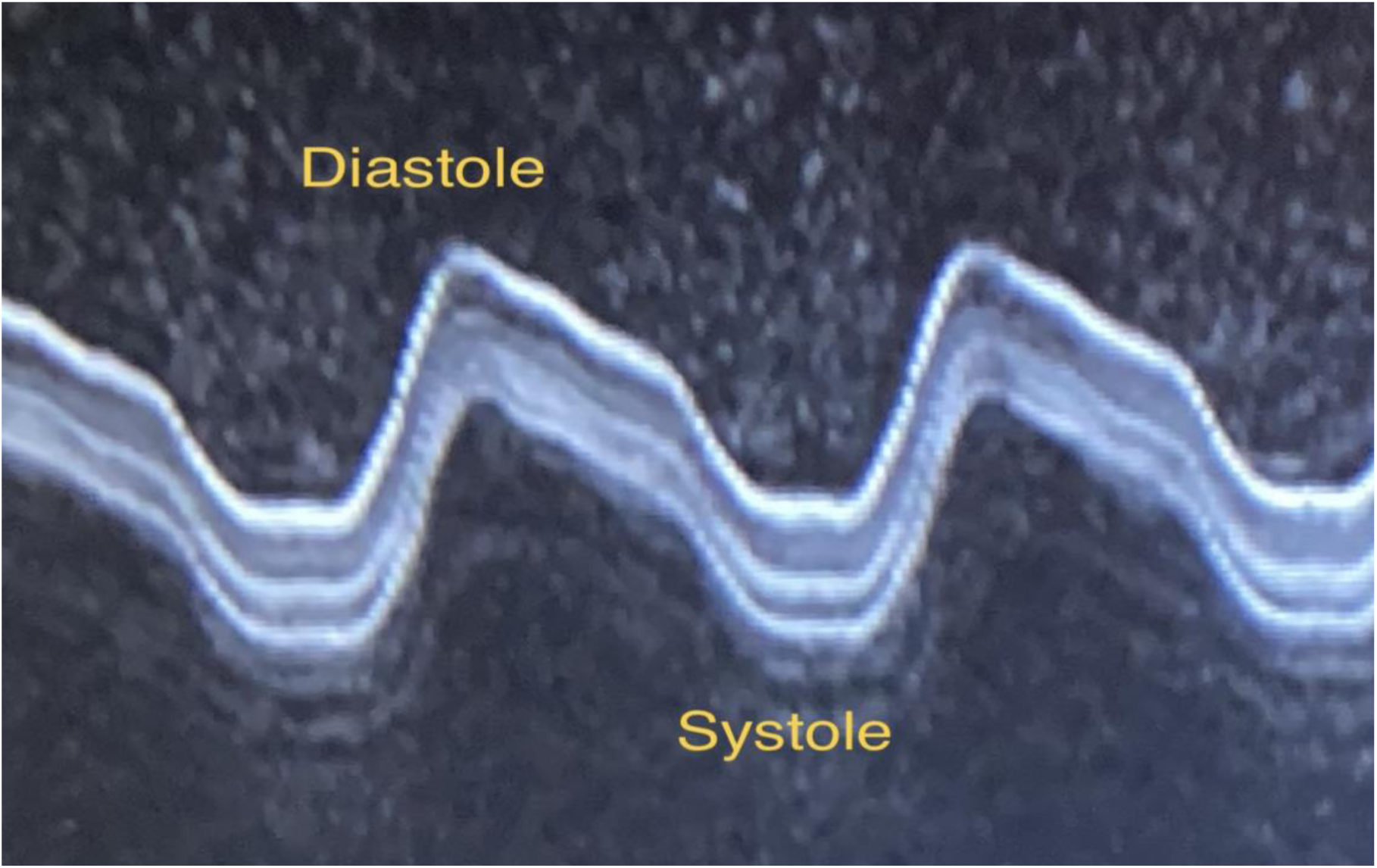
M-mode ultrasound showing differential strain separation of the layers of the posterior porcine aortic wall including the internal and external elastic lamina during passage of the pulse wave. The echoes below the arterial wall are reverberation artefacts in the water bath surrounding the artery. Peak layer separation is seen over the systolic/diastolic and diastolic/systolic transitions.

### Arc stretching

When the mounting tubes were approximated to each other to allow a degree of longitudinal kinking of the artery with the passage of each pulse wave, this kinking was seen to cause the plaque to lift on its rigid baseplate and to intrude into the vessel lumen during longitudinal bending, and the surface area of the baseplate in direct contact with the magnet across the arterial wall was consequently diminished according to the degree of bend. The sharper the bend the greater the CDF necessary to retain the plaque (Rs 1; n =5), Fig. 5.

**Figure 5.**
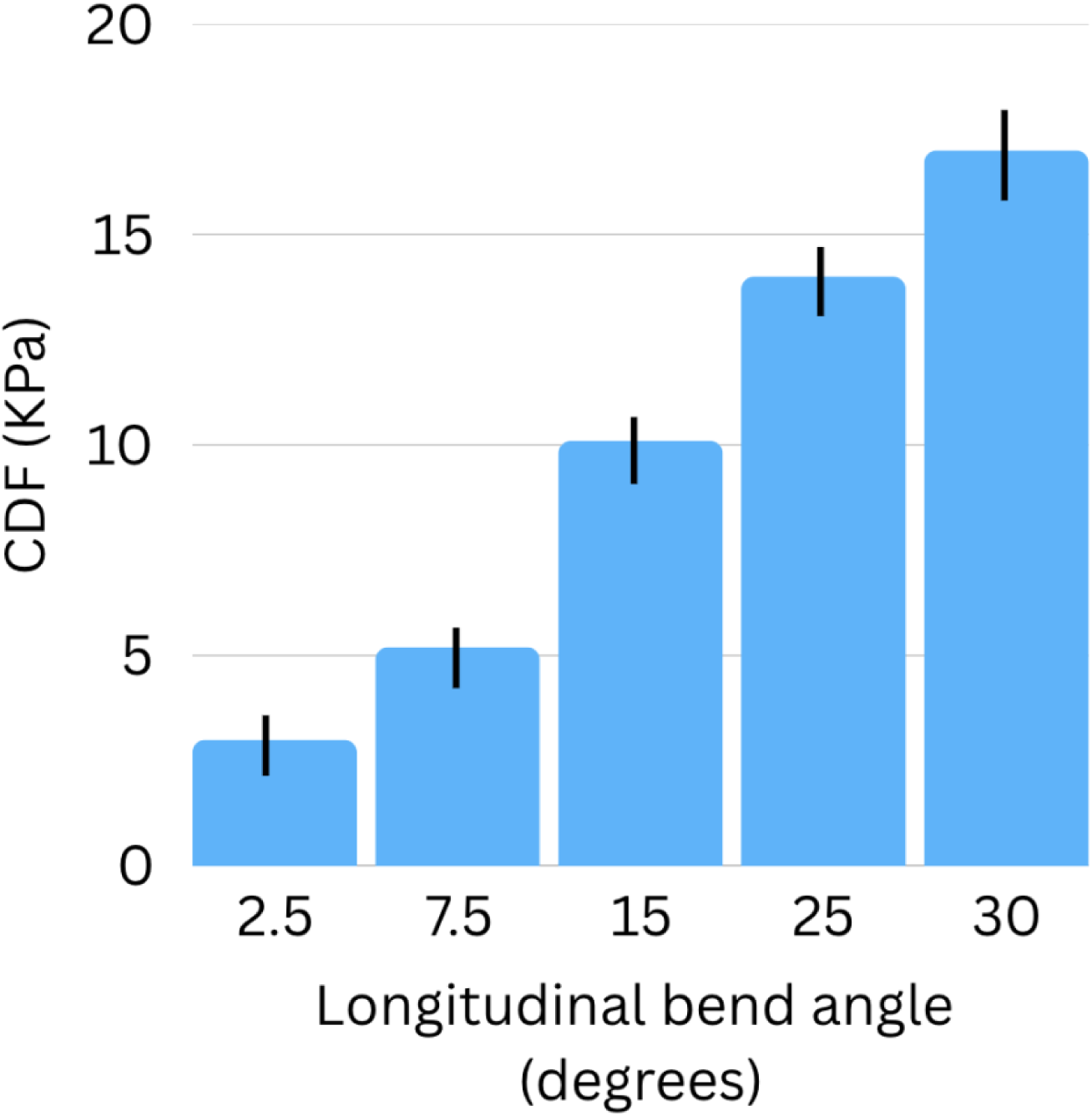
The effect of kink angle on CDF. The ends of the vessel have been approximated by stages so that its form becomes curved the artificial plaque being positioned on the greater curvature of the bend. As the pulse wave passes the vessel flexes in the longitudinal plane with the relatively rigid plaque being projected towards the lumen during diastole and losing its contact with the underlying wall.

## Discussion

The vulnerable atheromatous plaque has been identified on the basis of imaging studies and histology in specimens from acute coronary syndrome patients, as one with a large soft lipid core, capsular thinning, underlying haemorrhage or neo-vascularisation, stress concentration due to calcification, and the presence of an inflammatory infiltrate (Box A) . Less is known about the haemodynamic contribution to plaque rupture, and in particular about the interplay of dynamic factors, although ultimately it is mechanical stress is that separates the plaque from the vessel wall or causes capsular rupture. ^9^

**Box A. Recognised plaque-related factors contributing to plaque vulnerability**

|  |  |  |
| --- | --- | --- |
| Plaque | Spotty calcification<br>Low attenuation soft lipid core<br>Inflammation; napkin ring sign<br>Capsular thickness<br>Profile<br>Intrinsic tension | Hutcheson et al 2014 <sup>20</sup><br>Falk E et al 1992 <sup>2</sup><br>Otsuka K et al 2013 <sup>24</sup>; Moreno et al 1994 <sup>25</sup> Shah 1997 <sup>4</sup><br>Virmani et al 2003 <sup>26</sup><br>Finet et al 2007 <sup>23</sup><br>Ohayon et al 2007 <sup>27</sup> |
| Flow | Pulse pressure | Li et al 2020 <sup>28</sup>; Glyn et al 2000 <sup>29</sup> |
| Underlying wall | Compliance<br>Bending/<br>Branching | Rezvari-Sharif et al 2018<sup>30</sup><br>Finet et al 2007 <sup>23</sup><br>Yang et al 2008 <sup>21</sup> |
| Rheological | Hyper-viscosity | Koenig et al 1998 <sup>19</sup> |
| Stenosis | Degree of stenosis | Falk et al 1995 <sup>5</sup>;<br>Shah 1997 <sup>4</sup> |

In principle there are three possible sources of haemodynamic stress acting on the plaque:

1. the force acting centripetally tending to lift the plaque off the vessel wall, due to the Venturi /Bernouilli effect. This force is present in an expansile vessel even in the absence of plaque, and under these circumstances reaches its maximum at the systolic/diastolic transition as the vessel narrows but flow velocity initially remains high. ^1^ The magnitude of the force is related to peak flow velocity, the form of the pulse pressure wave (whether rounded or peaked) and the degree of occlusion and thereby indirectly to pulse pressure. This may account for the tendency of a soft plaque to rupture over the dome of the plaque when subjected to a high pulse pressure. ^9,10^ It may also account for the progressive long-term stiffening of large central arteries through a mechanism involving repetitive stress>inflammatory change>wall remodelling ^(^Fig. 6). ^11–13^

**Figure 6.**
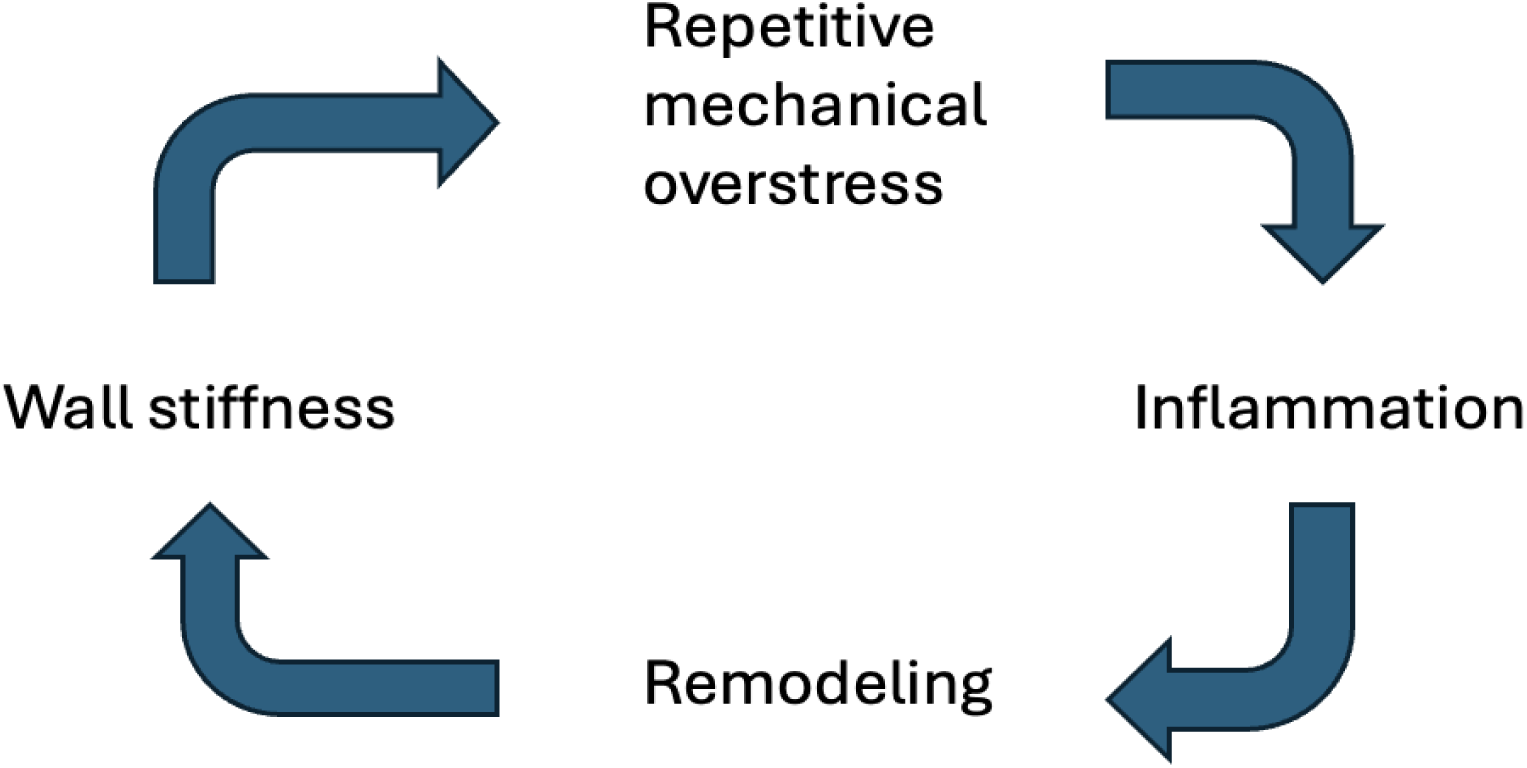
The proposed vicious circle of repetitive strain, inflammation, wall remodelling and large artery wall stiffness.

The hypothesis that repeated dynamic stress is responsible for inflammatory change in the arterial wall is supported by the link observed between arterial wall stiffness and elevated C-reactive protein (CRP), a marker of the systemic inflammatory response ^14^, and the observation that in hypertensive patients with elevated CRP, the CRP reduces in response to a variety of antihypertensive treatment and consequently to a reduction in arterial wall stress. ^15^

2. The torsional component of wall shear stress. This tends to dissect the plaque off the wall at the upstream shoulder region, where the majority of plaque ruptures are found to occur in post mortem findings in acute coronary artery occlusion victims, and where intravascular imaging demonstrate high stress and strain. ^16–18^ The effect of shear stress accounts for the role of viscosity noted both in these experiments and epidemiological studies in relation to cardiovascular risk. ^19^
3. A shear force acting between the plaque and the underlying wall caused by the disparity between the elastic qualities of the wall and those of the plaque during cyclic bending and circumferential stretch; in the current experiments it notably occurs in relation to longitudinal arc bending. Clinically it is recognised by the contribution of arc bending and of focal calcification, with local hardening of the plaque, to plaque vulnerability. ^20,21^

The clinical relevance of these observations is that pulse pressure, waveform profile, plasma viscosity and wall stiffness are all potentially modifiable, and so the risk of plaque rupture could be reduced in at-risk cases by targeting these elements therapeutically. Furthermore the recognition of the dual role of plaque-related factors (stenosis, inflammation, haemorrhage and compliance mismatch) and patient-related factors (pulse pressure, viscosity, wall stiffness) in determining plaque vulnerability provides a route for developing a scoring system based on non-invasive imaging and clinical criteria such as pulse pressure and CRP to define the level of risk, and to predict the most effective clinical strategy for reducing it. ^22^

These experiments have significant limitations: though the in-vitro fresh artery model can provide evidence about the main dynamic elements contributing to plaque separation, the arteries used were large elastic arteries, rather than coronary arteries, and the artificial plaques were composed of stiff epoxy resin bound to thin metal strips. The sucrose solutions used to test the effect of increased perfusate viscosity lacked the shear-thinning properties of blood. Furthermore, the model used fresh arteries outside the context of a living system and was therefore unable to explore the mechanism of the postulated link between recurrent mechanical stress and both local and systemic inflammatory effects. ^22^ These effects might account for the local accumulations of macrophages around the plaque at points of dynamic stress, ^18,23,25,31^ and would provide an explanation for the previously-mentioned association between inflammatory indicators such as CRP with cardiac risk, and the reversal of this association by the anti-inflammatory effect of statins and anti-hypertensives. The idea that the inflammatory component of plaque vulnerability is the result of a repetition strain injury acting on points of stress between the plaque and its attachments would be further supported if one could demonstrate that such repetitive strain gives rise to the expression of signalling molecules causing endothelial-macrophage adherence and translocation. A candidate molecule is the elastin degradation peptide-elastin receptor complex. ^32^ This has been shown to have such signalling effects in patients with susceptible elastic tissue due to Marfan’s disease or proximal thoracic aortic aneurysm. ^33^ It has also been established that static stress acting on elastic tissue in arteries will speed up elastin degradation in the presence of elastase; ^34^ the missing link is the demonstration that recurrent stress acting on the elastic tissue in the vessel wall would give rise to such signalling in the absence of prior enzymatic degradation or abnormal phenotype. This might be a fruitful area for further research with a view to developing agents other than statins that could block the inflammatory consequences of mechanically-induced elastin degradation at stress points around the plaque.

Figs. 7 proposes a summary of this integrated view of the proposed direct and indirect role of biomechanical strain in plaque rupture and emphasizes the importance of central vessel arterial wall stiffness in determining pulse pressure and transmural strain.

**Figure 7.**
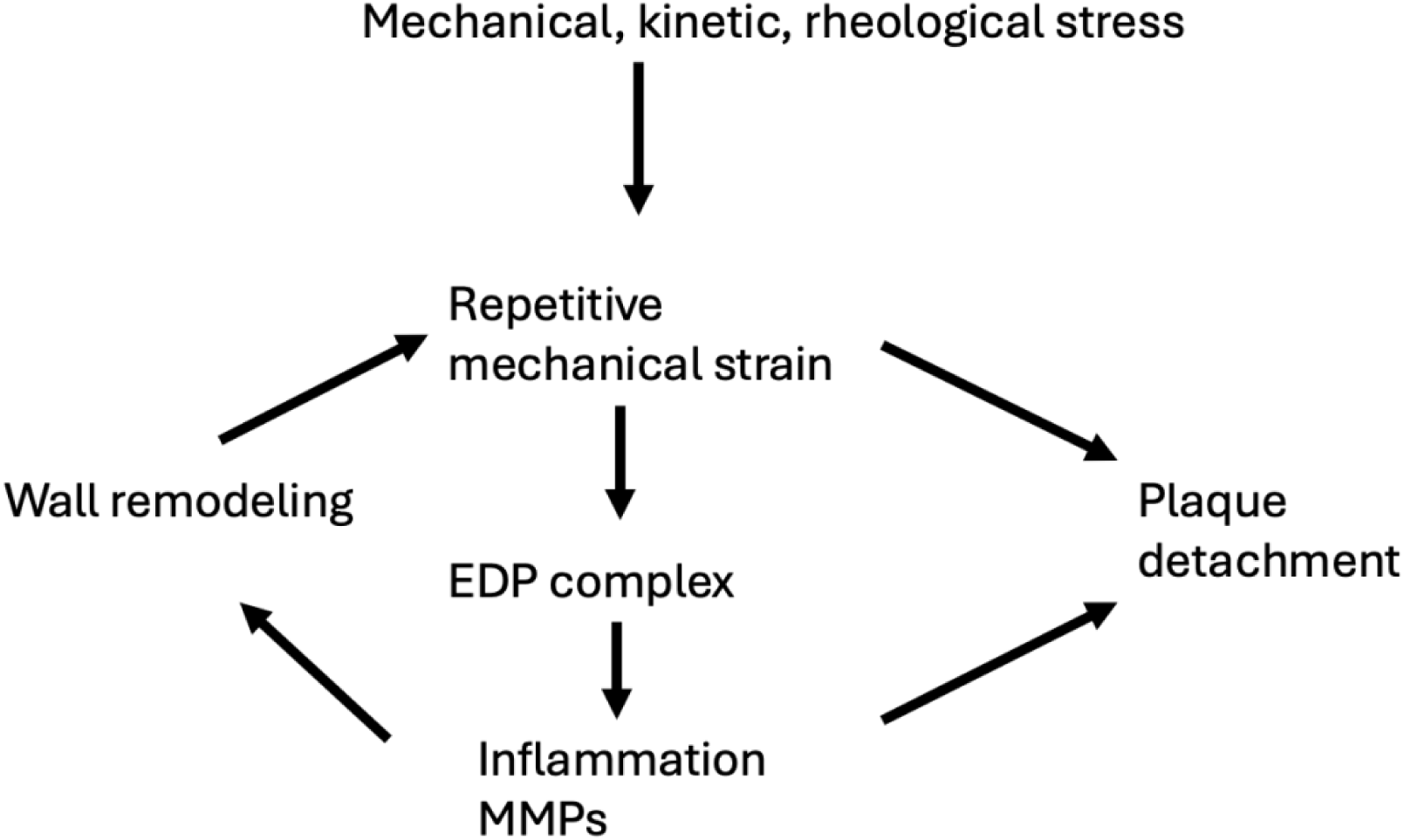
An integrated interpretation of the interaction of mechanical forces and inflammatory changes contributing to arteriosclerotic plaque detachment. EDP = elastic degradation peptide. MMP = matrix metalloproteinase.

## Conclusions

The mechanical forces tending to disrupt an artificial plaque relate to the form and the magnitude of the pulse pressure wave. These in turn depend on arterial wall stiffness within the proximal central distributive vessels. Viscosity and longitudinal bending also play a part, an area of further investigation. These findings are coherent with what is known about risk factors for acute adverse cardiovascular events.

Mechanical stress appears to act not only directly on the plaque but also indirectly by causing inflammatory changes. Histopathology and imaging studies have previously shown focal infiltration of macrophages and release of lytic enzymes at points of stress concentration where the plaque is prone to rupture. This inflammatory change is reflected in a rise in CRP which subsequently tends to fall if the mechanical stress or its inflammatory consequences are reduced with anti-hypertensive drugs or statins. In this context sensitive measurement of CRP (hs -CRP) is likely prove to be a useful gauge of risk, and of the effectiveness of antihypertensive or statin treatment, irrespective of cholesterol levels. It may also help to distinguish those younger hypertensive patients who would benefit from anti-hypertensive treatment from those with “white coat” hypertension, and it is proposed that patients at risk with consistently elevated highly sensitive CRP should be considered for statin treatment irrespective of their cholesterol status. The mechanism underlying plaque separation implicates the sequence *repetitive strain>inflammation>plaque rupture*. Measures to reduce the strain, for example by reducing elevated pulse pressure, by improving central vessel wall elastic compliance might be expected to reduce that risk. It is also important to explore the link between repetitive mechanical strain and inflammation. It is unfortunate that, although richly endowed with sympathetic nerves, large arteries themselves do not hurt when repeatedly strained. A device that could warn us when our pulse pressure exceeds a certain value might save lives.

## Acknowledgements

The authors would like to thank M. Joël Craveur for his invaluable technical assistance.

